# How do citizens perceive farm animal welfare conditions in Brazil?

**DOI:** 10.1101/380550

**Authors:** Ricardo Guimarães de Queiroz, Carla Heloisa de Faria Domingues, Maria Eugênia Andrighetto Canozzi, Rodrigo Garófallo Garcia, Clandio Favarini Ruviaro, Júlio Otávio Jardim Barcellos, João Augusto Rossi Borges

**Author notes:** Corresponding author, (JARB). These authors contributed equally to this work (Ricardo Guimarães de Queiroz, Carla Heloisa de Faria Domingues, Maria Eugênia Andrighetto Canozzi, João Augusto Rossi Borges). These authors also contributed equally to this work (Rodrigo Garófallo Garcia, Clandio Favarini Ruviaro, Júlio Otávio Jardim Barcellos).

## Abstract

The aim of this study is to understand the perceptions of Brazilian citizens about the actual conditions of farm animal welfare in the poultry, beef, and dairy supply chains. To reach this aim, an online survey was conducted. The analysis was based on descriptive statistics and three logistic regressions models. Results of descriptive statistics showed that citizens in Brazil had mostly negative perceptions about the actual conditions of animal welfare in the poultry, beef, and dairy supply chains. Results of the logistic regression models showed that in the poultry and dairy supply chains, citizens with background in agricultural/veterinary sciences, and citizens who reported a higher level of knowledge about these supply chains, were more likely to perceive as bad the actual conditions of farm animal welfare. In the poultry supply chain, citizens who reported previous contact with poultry farms were also more likely to perceive as bad the actual condition of farm animal welfare. In addition, the perception that farmers are mainly focused on the economic aspect of farming and less on animal welfare, the perception that animals do not have a good quality of life while housed on farms, and the perception that animals are not adequately transported and slaughtered, negatively impact on perceptions about the actual conditions of farm animal welfare in the three supply chains. We concluded that a protocol aimed to improve citizens’ perceptions about the actual conditions of farm animal welfare should focus in all phases of the supply chains.

## Introduction

In the last decades, there is an increasing public concern about the welfare of animals used for food production, with citizens, particularly from developed countries, questioning the intensification of animal production systems and requiring that farm animals have a good life [1]. In some countries, pressure from society has led to changes in animal production systems, which resulted in improvements of farm animal welfare (FAW) standards [1 – 3]. However, in some cases, changes in animal production systems may not be well suited for all stakeholders in food supply chains [4]. Therefore, if we want to successfully implement strategies to improve FAW standards, it is important to understand the concerns and perceptions of all stakeholders involved in food supply chains [1]. Particularly important is the understanding of society’ perceptions about FAW, because citizens play an important role in determining what is acceptable or not when it comes to FAW. For instance, citizens can pressure the government to implement laws to improve the welfare of animals used for food production or they can choose to buy certain type of products that guarantee good FAW standards [5].

A common approach used by researchers to investigate societal attitudes and perceptions is to carry on surveys about a specific farming practices or animal production systems that impact on FAW [4, 6–11]. Another approach is to carry on surveys to investigate more general perceptions (e.g. awareness) about FAW [12–16]. Most of these studies are conducted in developed countries in Europe and North America. In Brazil, one of the leading countries in livestock production, studies on society perceptions about FAW are emerging but there is a need to deeper our understanding of how Brazilian society perceives FAW conditions. Previous research conducted in Brazil has focused on citizens’ perceptions about specific farming practices [3, 17] and animal production systems that impact on FAW [18]. Our study moves beyond the previous literature by focusing on Brazilian citizens’ perceptions about the actual conditions of FAW on poultry, beef, and dairy supply chains and in the factors that might explain their perceptions. These factors include socio-demographic characteristics, awareness about animal welfare, knowledge about supply chains, perceptions about farming, perceptions about the quality of life of farm animals, perceptions about the use of animals for human consumption and perceptions about the conditions of transport and slaughtering in each supply chain. Therefore, the aim of this was to understand the perceptions of Brazilian citizens about the actual conditions of farm animal welfare in the poultry, beef, and dairy supply chains.

## Material and methods

### Survey and sampling

We developed three similar questionnaires: one for poultry supply chain, one for beef supply chain, and one for dairy supply chain. The questionnaires consisted of three groups of questions. In the first group, we measured participants’ socio-demographic characteristics. In the second group, we measured participants’ perceptions about the actual conditions of FAW on each supply chain and other questions related to animal welfare. All the variables and scales used are presented in S1 Table. In the third group, we used statements to measure participants’ perceptions about animal welfare. The statements were adapted from Boogaard et al. [12]. Statements used to measure participants’ perceptions are presented in S2 Table. All questions and statements were specifically adapted for each of the three supply chains. This project received research ethics board approval from Federal University of Grande Dourados/Faculty of Management, Accounting and Economics. Before starting data collection, the questionnaire was tested with 20 participants. All the questions were translated to Portuguese.

To collect the data, we conducted an anonymous online survey. In a first step, we contacted by phone human resource departments in several universities across Brazil. In this first contact, we explained the purpose of our research, and asked if the department would forward a survey link for the personal e-mail of students, professors and administration staff. Upon acceptance, we sent a follow-up e-mail to human resource departments with the survey link and a brief description of the research, which was then disseminated online for the academic community. Each university disseminated the questionnaire of only one supply chain. We received 1.617 questionnaires of which three were disregarded because they were incomplete. The final number of questionnaires was 728 for the poultry supply chain, 586 for the beef supply chain, and 300 for the dairy supply chain. The data collection took place from November 2016 until December 2017.

### Statistical analysis

Statistical analysis was conducted in two steps. In a first step, we used factor analysis to reduce the number of items used to represent participants’ perceptions about animal welfare. Principal component was used as the extraction method. The criterion to define the number of factors was an eigenvalue greater than one [19]. Items were included in a factor when they presented factor loadings greater than 0.5. Factors scores were generated for subsequent analysis [19].

In a second step, we run three logistic regression models. The three dependent variables were participants’ perceptions about the actual conditions of FAW on each supply chain. In the original questionnaires, this variable was measured in a Likert scale from 1 to 5 (S1 Table). In order to run the logistic models, we transformed the variable participants’ perceptions about the actual conditions of FAW on each supply chain into a binary variable, where participants who answered 1 or 2 were gathered to a bad condition group (Bad:0) and participants who answered 3, 4 or 5 were gathered to regular condition group (Regular:1). We tested the impact of two groups of independent variables: participants’ socio-demographic characteristics, and participants’ perceptions about animal welfare. The significance level was p<0.05.

## Results

### Descriptive statistics

Descriptive statistics of participants’ socio-demographic characteristics are presented in S3 Table. Socio-demographic characteristics were similar for the participants in the poultry and beef supply chains, but somehow different for participants in the dairy supply chain. Participants who answered the dairy supply chain questionnaire were, on average, older, more educated, and earned a higher income compared to participants who answered the poultry and beef supply chains questionnaires. Apart from these differences, others participants’ socio-demographic characteristics were similar within the three supply chains: most participants were female, most of them study/work out of the fields related to agricultural/veterinary sciences, most of them had previous contact with farm animals, most of them lived in urban areas, and most of them were pet owners.

Descriptive statistics of participants’ perceptions about the actual conditions of FAW on each of the three supply chains and other questions related to animal welfare are presented in S3 Table. In general, participants within the three supply chains were aware of the concept of animal welfare. However, most participants reported that they do not have a high level of knowledge about animal welfare regulations. Within each of three supply chains, participants reported a medium level of knowledge about the supply chains. Within each of three supply chains, most participants perceived the actual conditions of FAW as very bad, bad, or regular, and few participants perceived the actual conditions as good or very good. In the poultry and beef supply chains, most participants did not agree that chickens and cattle are transported and slaughtered adequately.

### Factor analysis

Results of the factor analysis are presented in S4 Table. For the three supply chains, there were three factors with eigenvalue above 1.0. These three factors explained 67.5%, 62.1%, and 63.7% of the total variance in the poultry, beef, and dairy supply chains, respectively. Results of the factors loadings were also similar in the three supply chains. Following Boogaard et al. [12], we named the first factor ‘Farmers’ Image (FI)’, the second factor ‘Life Quality of Farm Animals (LQ)’, and the third factor ‘Use of Animals for Human Consumption (HC)’. The first factor describes participants’ perceptions about farmers. The items of this factor were negatively formulated in the questionnaire, so the higher participants scored on FI, the more they agreed that farmers are mainly focused on the economic aspect of farming and less in animal welfare. The second factor describes participants’ perceptions of the actual conditions of animal welfare in farming. The higher participants scored on LQ, the more they agreed that animals have a good quality of life while housed on farms. The third factor describes participants’ perceptions about the use of animals for human consumption. The lower participants scored on HC, the more they agreed that humans are allowed to use animals for consumption.

Descriptive statistics about the statements used to measure participants’ perceptions about animal welfare are presented in S2 Table. For the statements related to FI (Perc_1_, Perc_2_, Perc_3_, Perc_4_), the mean were above or close to 4, which indicates that participants agreed that most farmers focus too much on the economic aspect of farming and less in animal welfare. For the statements related to LQ (Perc_5_, Perc_6_, Perc_7_, Perc_8_), the mean were below or close to 3, which indicates that participants did not agree that animals have a good quality of life while housed on farms. For the statements related to HC (Perc_9_, Perc_10_), the mean were below or close to 2, which indicates that participants agreed that humans are allowed to use animals for consumption.

### Logistic regression models

We tested the impact of socio-demographic characteristics, and participants’ perceptions about animal welfare on their perceptions about the actual condition of FAW in each supply chain. Results of the three logistic regression models are present in Table 1. The socio-demographic characteristics age, gender, pet ownership, and consumption of animal products did not significantly impact on participants’ perceptions about the actual condition of FAW in any supply chain. In the poultry supply chain, participants who reported previous contact with poultry farms were more likely to perceive as bad the actual condition of FAW compared to participants who had not reported previous contact. In the poultry and dairy supply chains, participants in the fields of study related to agricultural/veterinary sciences were more likely to perceive as bad the actual conditions of FAW compared to participants out of these fields. In those supply chains, participants who reported a higher level of knowledge about poultry and dairy supply chains were more likely to perceive as bad the actual conditions of FAW compared to those participants who reported a lower level of knowledge about these supply chains. Participants’ perceptions about animal welfare also influence their perceptions about the actual conditions of FAW. In the poultry and beef supply chains, participants who perceived that animals are adequately transported and slaughtered were more likely to perceive as regular the actual conditions of FAW compared to those participants who perceived that animals are not adequately transported and slaughtered. Within each of three supply chains, participants who perceived that farmers are mainly focused on the economic aspect of farming and less in animal welfare (FI) were more likely to perceive as bad the actual conditions of FAW compared to those who perceived that farmers are more focused on animal welfare and less in the economic aspect of farming. Moreover, participants who perceived that animals have a good quality of life while housed on farms (LQ) were more likely to perceive as regular the actual conditions of FAW compared to those who perceived that animals do not have a good quality of life while housed on farms. Finally, participants who perceived that humans are allowed to use animals for consumption (HC) were more likely to perceive as regular the actual conditions of FAW compared to those who perceived that humans are not allowed to use animals for consumption.

**Table 1.**
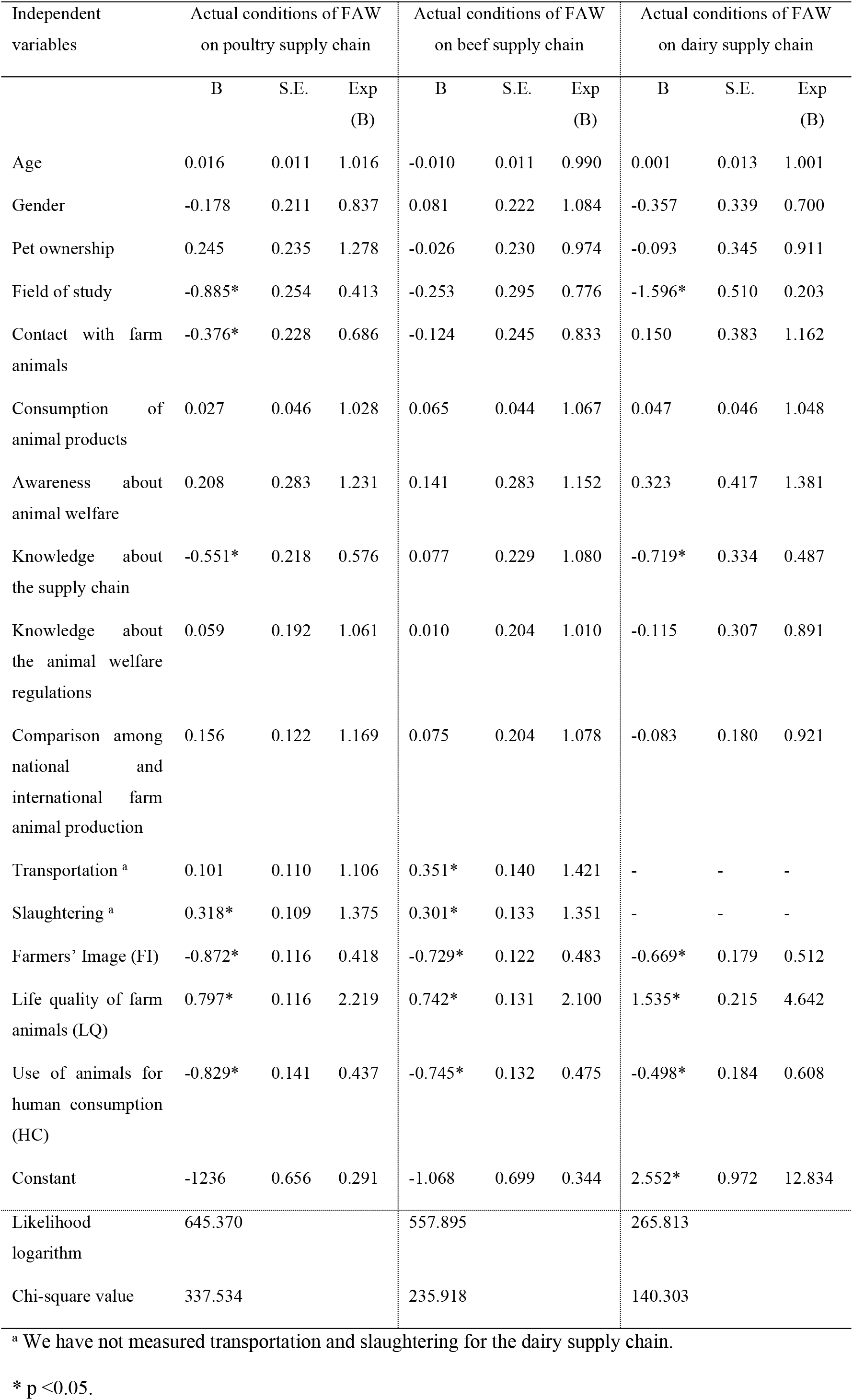
Logistic regression models of the Brazilian citizen perceptions about the actual conditions of FAW on poultry, beef, and dairy supply chains

## Discussion and concluding comments

Using a general measure of the perceptions about the actual conditions of FAW, we found that the vast majority of participants perceived the actual conditions of FAW in the poultry, beef and dairy supply chains as very bad, bad, or regular. When we specifically asked participants about their perceptions about the animal welfare conditions in farming, most of them did not agree that farm animals have a good quality of life. In their review, Clark et al. [20] showed that citizens from developing and developed countries have more negative than positive perceptions about farm animal welfare conditions, which is in line with our findings. Our results also showed that most participants agreed that farmers focus much more on economic aspects of farming and less on animal welfare, and most of the participants did not agree that animals are adequately transported and slaughtered. In summary, most participants in our sample had general and specific (farming, transportation, and slaughtering) negative perceptions about the actual conditions of FAW in the poultry, beef, and dairy supply chains in Brazil.

Results of the three logistic regressions were similar. Most socio-demographic characteristics did not impact on perceptions about the actual condition of FAW in any of the three supply chains. In contrast, Kupsala et al. [13] found that women, younger people, and people who are pet owners perceived the actual conditions of FAW in Finland more negatively than men, older people and people who are not pet owners. These contradictory results might be explained because, while Kupsala et al. [13] focused on general public, our sample is restricted to academic community, where socio-demographic characteristics play a lesser role in explaining the variation in perceptions about the actual condition of FAW. We recommend that future research focus on Brazilian general public to investigate the role of socio-demographic characteristics in shaping perceptions about FAW conditions.

In our logistic regression models we had three variables related to participants’ knowledge about the supply chains: their background in agricultural/veterinary sciences, a self-reported level of knowledge and previous contact with farms. Results of the logistic regressions models showed that these variables related to knowledge about the supply chains negatively impact on perceptions about the actual conditions of FAW in the poultry and dairy chains. These results can be explained by a growing body of literature indicating that as more people know about farming practices, the more they think that these practices do not provide a good quality of life for farm animals [9, 11, 17]. In contrast, results of the logistic regression model for the beef supply chain showed that variables related to knowledge about the supply chain and farming did not impact on perceptions about the actual conditions of FAW. These results might be explained by the difference in animal production systems used in the three supply chains in Brazil. The predominant production systems in poultry and dairy supply chains in Brazil are intensive, where animals live mostly confined [1, 17] whereas, in the beef supply chain, animals are reared in more extensive systems [21]. Intensive production systems are usually perceived by citizens as unnatural and by providing less FAW compared to extensive systems [20]. Therefore, in our sample, participants who had more knowledge about animal production systems in the poultry and dairy supply chains might know that mostly animals are housed in confinement housing systems, and were more likely to perceive the actual conditions of FAW in these two chains as bad. In contrast, in the beef supply chain participants who have more knowledge about animal production systems might know that mostly animals are reared in extensive production systems, and therefore knowledge did not impact on their perceptions about the actual conditions of FAW. These results suggest that increasing citizens’ education about animal production systems and practices used in supply chains will decrease their acceptance of such production systems and practices, particularly in supply chains with more intensive production systems. Ventura et al. [22] also claimed that education and exposure to livestock farming might not improve citizens’ perceptions that farm animals have a good life.

Results of the logistic regressions also showed that perceptions that farmers are mainly focused on the economic aspect of farming, perceptions that animals do have a good quality of life in farms, and perceptions that animals are not adequately transported and slaughtered, negatively impact on a general measure of the actual conditions of FAW. These results indicate that perceptions about animal welfare conditions on each phase of the supply chain shape the general perceptions about the actual conditions of FAW. Therefore, a protocol aimed to improve citizens’ perceptions about the actual conditions of FAW should focus in all phases of the food supply chains.

A potential limitation of this study concerns selecting participants only in the academic community. In comparison to the Brazilian population our sample is younger, more educated, and earns a higher income [23]. Although we acknowledge that our sample is unbalanced in terms of education, income, and age, we argue that academic community members have more access to information that might drive changes in production systems.

